# A conserved role of Parkinson-associated DJ-1 metabolites in sperm motility, mitosis, and embryonic development

**DOI:** 10.1101/2021.01.16.426934

**Authors:** Susanne Bour, Yanina Dening, Melanie Balbach, Ina Poser, Inés Ramírez Álvarez, Helena den Haan, Christoph Kluge, Ronald Naumann, Reinhard Oertel, Irene Alba-Alejandre, Davide Accardi, Christian G. Stief, Marianne Dieterich, Peter Falkai, Rainer A. Böckmann, Horacio Pérez-Sánchez, Anthony A. Hyman, Matthias Trottmann, Francisco Pan-Montojo

## Abstract

Fertility rates in the developing world have dramatically dropped in the last decades. This drop is likely due to a decline in sperm quality and women having children at older ages. Loss of function mutations in *DJ-1*, a Parkinson’s associated gene, are linked to alterations in multiple cellular processes such as mitochondrial activity, ROS production or sperm motility and lead to an early onset of Parkinson’s disease and male infertility in humans and other species. Glycolate (GA) and D-lactate (DL), products of *DJ-1* glyoxalase activity, sustain mitochondrial function and protect against environmental aggressions. We, therefore, tested whether these substances could also have a rescue effect on these phenotypes. Here, we show that *DJ-1* loss of function not only affects sperm motility but also leads to defects in mitosis and an age-dependent increase in the abortion rate. Remarkably, whereas DL was only able to rescue embryonic lethality in *C. elegans*, GA rescued these phenotypes in all model systems tested and even increased sperm motility in wild-type sperm. These positive effects seem to be mediated through an increase in NAD(P)H production and the regulation of intracellular calcium. These findings not only strongly suggest GA as a new therapeutic candidate to improve male and female fertility but also show its potential to treat diseases associated with a decline in mitochondrial function or to improve mitochondrial function in aging.

## Introduction

Fertility rates in the developing world have dramatically dropped in the last decades (reviewed in ^1^). This drop is likely due to a decline in sperm quality, specially a reduction in spermatozoa concentration and motility, and a shift in the mean age at which women decide to start having children. Alterations in sperm quality and increases in the abortion rate have been related to aging and to exposures to environmental toxins. Therefore, looking for treatments that could improve sperm quality and increase fertility protecting against environmental agressions is of the essence.

Loss of function mutations in *DJ-1* (also known as *PARK-7*) are associated with mitochondrial dysfunction leading to early onset of familial Parkinson’s disease (PD) ^2^. These mutations have also been associated with defects in sperm motility and male infertility in humans and other species ^3-5^. At the cellular level, *DJ-1* loss of function mutations are associated with abnormal mitochondrial morphology and dynamics, increased sensitivity to oxidative stress, opening of the mitochondrial permeability transition pore, and alterations in calcium homeostasis ^6-8^. Despite intensive research in the last decade to elucidate the molecular mechanisms underlying this plethora of cellular functions, they still remain unknown.

DJ-1 has recently been identified as a member of a novel glyoxalase familiy^8^. Two systems of glyoxalases have been described: 1) Glutathione-dependent Glo I and Glo II systems (GLOD-4) ^9^ and 2) cofactor-independent Glo III system (DJ-1) ^10^. Glyoxalases are enzymes that transform 2-oxoaldehydes (i.e. glyoxal and methylglyoxal) into their corresponding 2-hydroxyacids (i.e. glycolate (GA) and D-lactate (DL)). Glyoxal and methylglyoxal are potent aldehydes, covalently reacting with proteins and lipids to form advanced glycation end-products (AGEs), which have been implicated in PD and many other neurodegenerative diseases ^11,12^. Thus, DJ-1 and GLOD-4 were considered to act as detoxifying proteins. GA and DL were first described more than a century ago as a component of the sugar cane by Shorey ^13^ and a product of glycolysis and amino acid metabolism by Dakin and Dudley ^14^, respectively, but their function remained unknown for a long time. The results of our previous study suggested that these glyoxalases have two different but complementary functions: They detoxify harmful substances produced by glycolysis (i.e. glyoxal and methylglyoxal) and they produce GA and DL, two substances that support mitochondrial function. We have shown that knocking down *DJ-1* in HeLa cells and *C. elegans* leads to alterations in the mitochondrial membrane potential and increased sensitivity to oxidative stress caused by environmental toxins such as paraquat (PQ, a widely used pesticide associated with the appearance of PD) or desiccation ^15^. Interestingly, these phenotypes were rescued by the co-treatment with the products of *DJ-1*, GA and DL, in a dose-dependent manner. Moreover, the addition of these substances to dopaminergic neurons *in vitro* had a neuroprotective effect against PQ.

Low levels of *DJ-1* or its deletion have been associated with male infertility in humans, flies and mice, mostly due to defects in sperm motility ^3-5^. In this paper we show that knocking down *djr-1*.*1/djr-1*.*2* or *glod-4* in worms and *DJ-1* in mice did induced a reduction in the brood size of worms and in the litter size in mice. Interestingly, this reduction was not only due to a decrease in sperm motility but also to an age-dependent increase in the abortion rate, which is normally due to alterations in embryonic development. Treatment with GA or DL in worms and with GA, but only partially with DL, rescued these phenotypes in HeLa cells, sperm and worms and increased sperm motility even in wild type (WT) sperm.

## Materials & Methods

### Chemicals

HBSS without Ca^2+^ and Mg^2+^, PBS, poly-L-lysine, poly-D-lysine, CaCl^2^, glycolic acid, D-lactate, HEPES, ionomycin, Glutamic acid, hydrochloric acid solution, glucose (all from Sigma Aldrich, Germany, EU), DNAse I (Roche, Germany, EU). High glucose Dulbecco’s Modified Eagle Medium (DMEM, 31966-021), FBS, Pen/Streptomycin, trypsin, Fluor 4-AM, TMRE and Rhod-2 AM (all from ThermoFischer Scientific, USA). Anhydrous DMSO and Pluronic F-127 (Sigma Aldrich, Germany) were diluted at a 20% (w/v) and used to reconstitute Fluo-4 AM stocks right before use.

### Ethical statement

All animal procedures were performed according to the German Law for Animal Experiments (Tierschutzversuchsordnung). The ethical committee from the Munich University Hospital approved all procedures on human sperm (Az: 505-14).

### Determination of mouse litter size and abortion rate

To determine the abortion rate 8 to 30 weeks-old C57Bl6/J and PARK-7^-/-^ mice (n=16, matching ages) were sacrificed between day E14.5 and E15.5 and the number of embryos and abortions was determined. On a second batch of animals, litter size was determined as the number of newborn mice per pregnant mice (n=15).

### Determination of embryonic lethality and brood size

All *C. elegans* strains were maintained on NGM agar plates seeded with *Escherichia coli* NA22 at 15°C. Wild type (N2) and mutant strains *ΔΔdjr* and *glod-4*(tm1266) were obtained from Prof. Kurzchalia’s laboratory at the Max Planck Institute for Cell Biology and Genetics. The procedures to obtain the *DJ-1* double mutant mice have been already described ^15^. To determine embryonic lethality, individual adult worms from each strain were transferred to a 6-well plate well with NGM and *E. coli* (NA22) (with or without GA or DL) to lay eggs. After 4 hours, adult worms were removed and the number of laid eggs was counted. The percentage of hatched eggs was calculated (L1/(L1+remaining eggs)*100) 8 hours after removing the adults. To determine brood size, L4 worms were transferred to a 6-well plate well with NGM and NA22 and allowed to lay eggs. To prevent confusion, adult worms were transferred to another well every 24 hours until no laid eggs could be observed. The number of L4 and adult worms in each well were counted 24 hours after the adult worm had been transfered, and the brood size was determined for each strain.

### Determination of cell growth

Cell growth was determined by two different methods. The first method (WST1-Assay) was used to analyze cell growth at different time points using the same plates: 500 cells of 8 different PARK7 KO clones and HeLa Kyoto wild type cells were seeded in 96 well plates (6 wells / line). For each time point (0 h, 48 h, 122 h, and 144 h), WST1 was added to the cells according to the manufacturer’s instructions and incubated for 30 min at 37°C. Absorbance was measured at 450 nm and 620 nm using an EnVision Plate Reader (PerkinElmer).

The second method was used to analyze the rescue effect of GA and DL. Briefly, HeLa cells were seeded and treated with medium containing distilled water, 5 mM GA, or 5 mM DL. 48 hours later, the number of living cells was calculated with the help of an automated cell counter (ThermoFischer, USA).

### CRISPR

HeLa-Kyoto PARK7 KO clones had been kindly provided by Martin Stewart (Koch Institute, MIT, Cambridge, USA). Briefly, cells were electroporated with the NEON device (Invitrogen) using a sgRNA-Cas9-NLS complex targeting human PARK7 at exon 1 (sg1: CTCGCCTCATGACATCTACAGGG; sg2: AGATGTCATGAGGCGAGCTGGGG). Subsequently, cells were seeded in serial dilution and clones were characterized by genotyping, sequencing, and Western blot.

### Quantitative Western blot (anti PARK7)

Western blot was performed using anti-PARK7/DJ1 (sc-32874, SantaCruz) and anti-Tubulin (T9026, Sigma) antibodies and analyzed using an Odyssey imaging system (Licor).

### Semen sample collection and motility measurements

Mouse sperm samples were obtained from the epididymis of sacrificed C57Bl/6J or PARK-7-/-mice (n=3 per group). Briefly, both epididymis of each male were placed in 200 µl of an isotonic 0.9% NaCl solution and fat tissue was removed. Epididymis were then placed into 170 µl CPA buffer (18% raffinose and 3% milk powder in distilled water), cut into several pieces, and left at RT for 5 min to allow the spermatozoa to swim into the media. 4 µl of the sperm-containing suspension was then pipetted into 196 µl HTF medium supplemented with 10 mM of GA or DL or an equivalent amount of NaCl in distilled water to control for osmolality (vehicle) and incubated at 37°C for 60 min before starting the measurements. Mouse sperm motility measurements were performed using the Hamilton Thorne IVOS platform (Hamilton Thorne, USA) and the specific sperm motility analysis software for mice. Briefly, the IVOS platform was warmed up to 37°C before starting the measurements. 25 µl of sperm samples in HTF medium from each treatment group were loaded on a Leja-slide and introduced in the machine. After focusing on the samples, the machine analyzed 10 fields per samples. This analysis was performed before and 5, 30, 60 and 90 minutes after addition of vehicle, GA, or DL.

Frozen bull semen samples were provided from Swiss genetics, Switzerland. Human sperm samples were collected from patients enrolled in the study attending the diagnostic Department of the Urologic Clinic of the Ludwig-Maximilians University Munich for Semen Diagnostics and Artificial Insemination. All patients were between 28 and 58 years of age. Semen samples of 68 normozoospermic healthy patients were obtained by masturbation after 3 to 5 days of sexual abstinence. Some of the samples were diluted in SpermFreeze buffer (LifeGlobal, USA) and stored in liquid nitrogen for later analysis.

Fresh human samples were left at 37°C for 2 hours before semen samples were gently centrifuged (800 x g, 4 min) and resuspended in sperm iTALP buffer (99 mM NaCl, 3.1 mM KCl, 25 mM NaHCO_3_, 0.35 mM NaH_2_PO_4_ - H_2_O, 0.5 mM HEPES,2.0 mM CaCl_2_ - 2 H_2_O, 1.1 mM MgCl_2_ − 6 H_2_O and H_2_O dest, pH 7.4), containing 0.25 mg/ml Polivinylalcohol (PVA). On the day of analysis, frozen human and bull sperm samples were thawed at 37 °C, centrifuged (800 x g, 4 min), and the supernatant was exchanged with iTALP buffer pH 7.4. After re-suspension in iTALP buffer, human (fresh and thawed) and bull semen samples (n=10) were divided into treatment groups (vehicle (distilled water with NaCl), 30 mM or 60 mM GA with or without 1, 3, 5 and 10 µM of CatSper channel blocker NCC55-0396). Sperm motility was blindly and manually assessed following the WHO guidelines (2010) before treatment and 5, 30, 60, and 90 min after treatment. Mitochondrial activity was analysed by Fluorescent Activated Cell Sorter as described below.

### Fluorescence Activated Cell Sorting (FACS) of semen subpopulations

Semen samples were adjusted to 2.0 – 4.0 × 10^8^/ml and stained with JC-1 (2 µl/ml) in iTALP buffer for 10 min at RT. The samples were analyzed with a Becton Dickinson BD FACS Calibur Flow Cytometer. For detection we used the following settings: FSC (E00, 9.99 linear), SSC (510 mV, 1.00, linear), FL1 (480 mV, 1.00, logarithmic) with 488 nm wavelength and FL2 (520 mV, 1.00, logarithmic) with 543 nm wavelength. For each sample, 20,000 cells were counted. Compensation between two channels was adjusted to 59.9 %. Mitochondrial activity was evaluated by fluorescence emission of JC-1, which emits green fluorescence when the membrane potential is depolarized and red fluorescence when the membrane potential shifts to more negative values. Populations of cells were separated in depolarized and polarized mitochondrial membrane potential subpopulations by adjusting compensation and laser power in a way that two populations could be observed. For analysis, Flowing Software 2.0 (University of Turku, Finland) was used. Subpopulations were gated and analyzed accordingly.

### NAD(P)H live-cell microscopy on HeLa cells

NAD(P)H live-cell microscopy of HeLa cells was performed as previously described ^16^. Briefly, NAD(P)H fluorescence intensity time series were performed on a ZEISS LSM880 inverted confocal equipped with an incubation chamber to maintain 37 Celsius degree and 5% of CO2. Fluorophores were excited using a 355nm UV laser (Coherent), while the fluorescent signal was detected using a GaAsP spectral detector narrowing down the band of absorption between 455 and 473nm. To maximize the transmission efficiency of the system in excitation and detection and reduce the aberrations due to the watery environment, a ZEISS Plan C-ApoChromat 40x/1.2 Water lens with depth compensating correction collar was used. In addition, bright field images were taken using a HeNe 633 laser as source of light and a T-PMT to detect the signal. The sampling factor in XY (pixel size) of each image was equal to 208nm, which lead to a final resolution of approximately 600nm. For each image a volume of 5µm around the specimen central plane was taken by acquiring 3 planes separated by a Z-step of 2.5µm. Time series measurements were obtained with 5 min time resolution. Fluorescence intensity levels were extracted using FIJI Image Analysis Freeware

### Analyzing changes of the intracellular Ca^2+^ concentration in sperm and HeLa cells using a fluorescence plate reader

Mouse sperm were isolated by incision of the cauda epididymis followed by a swim-out in modified TYH medium (135 mM NaCl, 4.8 mM KCl, 2 mM CaCl^2^, 1.2 mM KH2PO4, 1 mM MgSO^4^, 5.6 mM glucose, 0.5 mM sodium pyruvate, 10 mM L-lactate, 10 mM HEPES, pH 7.4). After 15 min swim-out at 37°C, sperm were collected and counted. For capacitation, mouse sperm were subsequently incubated for 90 min in TYH containing 3 mg/ml BSA and 25 mM NaHCO_3_. Human sperm were purified by a “swim-up” procedure in human tubular fluid (HTF): 97.8 mM NaCl, 4.69 mM KCl, 0.2 mM MgSO_4_, 0.37 mM KH_2_PO_4_, 2.04 mM CaCl_2_, 0.33 mM Na-pyruvate, 21.4 mM lactic acid, 2.78 mM glucose, and 21 mM HEPES, pH 7.4. For capacitation, human sperm were incubated for 3 h in HTF containing 3 mg/ml human serum albumin and 25 mM NaHCO_3_. Changes in [Ca^2+^]_i_ were measured in 384-microtiter plates in a fluorescence plate reader (Fluostar Omega, BMG Labtech, Germany) at 30 °C. Mouse and human sperm were loaded with Cal-520-AM (5 μM, Molecular Probes, USA) in the presence of Pluronic F-127 (0.02% v/v) at 37°C for 45 min. After incubation, excess dye was removed by centrifugation (700 x g, 7 min, RT). Each well was filled with 50 µl of the sperm suspension (5·10^6^ sperm/ml); fluorescence was excited at 480 nm, emission was detected at 520 nm with bottom optics. After 10 cycles, the experiment was interrupted and 10 µl stimuli (mouse sperm: 10 mM GA/9 mM NaCl/10 mM 8-Br-cAMP, human sperm: 30 mM GA/27 mM NaCl/2 µM of progesterone) or buffer as control were added to the sperm suspension using an electronic multichannel pipette. After 60 cycles, 10 µl 2mM CaCl^2^ or buffer as control were added to each well.

Fluorescence measurements on HeLa cells were carried out in 96 well-plates (F-Bottom, Greiner, Germany, EU) using the fluorescence plate-reader Fluostar Omega (BMG Labtech, Ortenberg) with a 485/12 nm excitation filter with bottom optics for excitation and the Em520 nm filter for emission. 30.000 HeLa cells were plated in each well and left 12 hours at 37°C and 5% CO_2_ in the incubator overnight. On the next day, cells were loaded with Fluo-4 AM or TMRE as described for calcium imaging during mitosis. 20 min after the last HBBS change, cells were exposed to either 4,5 mM NaCl as osmotic control or GA (2,5 and 5 mM) after 5 min by adding 10 µl extra HBBS containing the substances (10x concentrated, 10% of the total volume) and 2 µM Ionomycin 20 min later.

## Results

### Knocking down *DJ-1* leads to a reduction of progressive sperm motility, an increased abortion rate and alterations in cell proliferation

First we investigated the effect of knocking down *DJ-1* in mice and *djr1*.*1/djr1*.*2* or *glod-4* in worms on litter and brood sizes. We used *DJ-1* KO (PARK-7^-/-^) mice, *djr-1*.*1;djr-1*.*2* double *C. elegans* mutants lacking both *DJ-1* orthologues *(ΔΔdjr*), and *glod-4 C. elegans* mutants lacking the Glo I/II system. Our results show that the loss of *DJ-1* in mice leads to a significant decrease in the litter size (Figure 1a). In worms, the loss of *djr-1*.*1* and *djr-1*.*2* and to a lesser extent the loss of *glod-4* leads to a significant reduction of the brood size (Figure 1b). To investigate whether this phenotype was due to alterations in sperm motility and/or embryonic development, we analyzed sperm motility in mice and the abortion rate/embryonic lethality in mice and worms respectively. As expected, we observed a reduction in the percentage of rapidly progressive motile sperm (Grade A motility according to the world health organization (WHO)) between sperm isolated from *DJ-1* KO mice and WT mice (Figure 1c). Interestingly and unexpected, we also observed an age-dependent increase in the abortion rate in *DJ-1* KO female mice compared to WT females (Figure 1d and 1e). This finding was also reproduced in *C. elegans*, where an increase in embryonic lethality in ΔΔ-*djr* and *glod-4* worms (measured as a reduction in the percentage of hatched eggs) compared to the control (Figure 1f) was observed.

**Figure 1.**
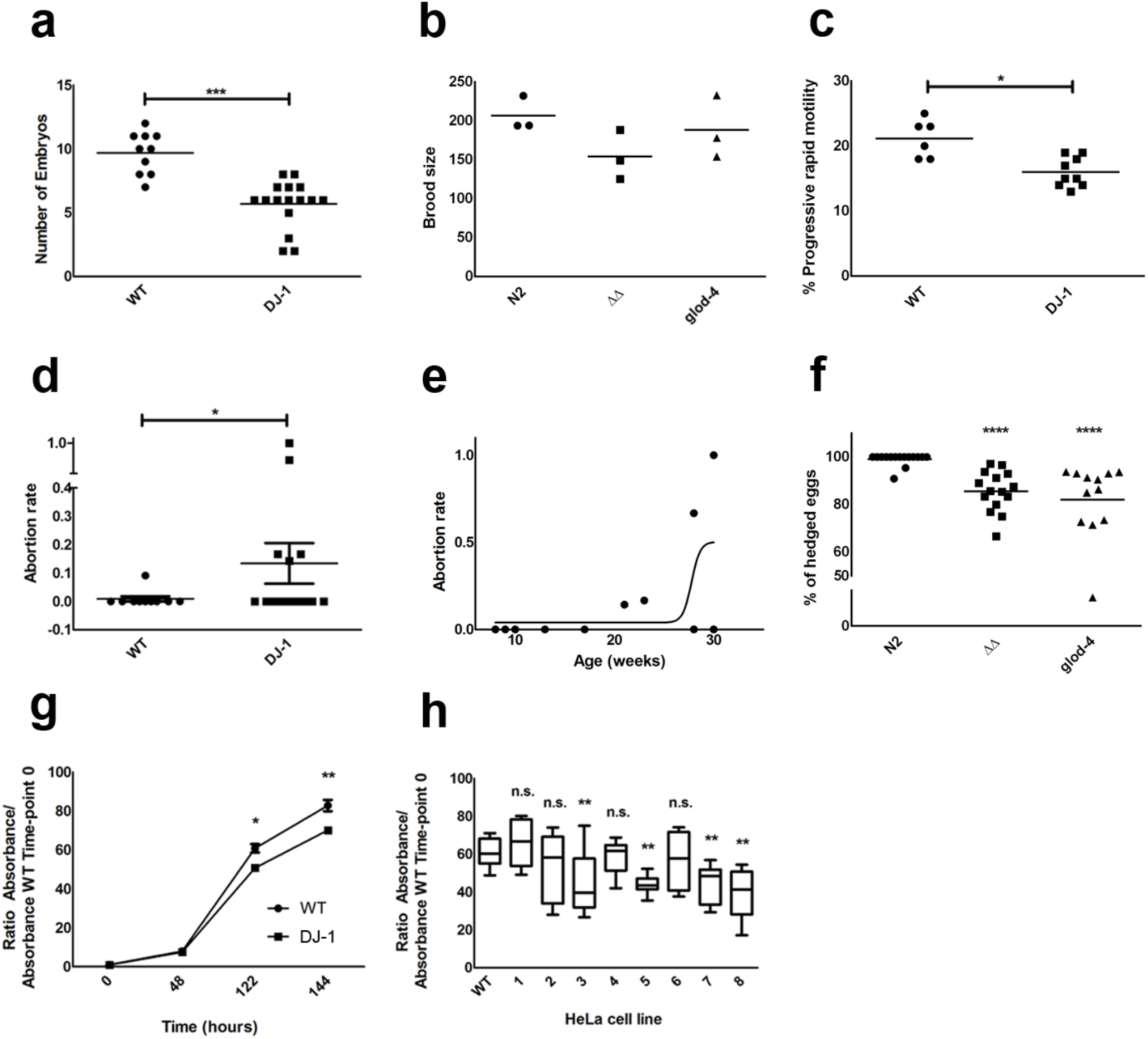
Effects of DJ-1 loss of function in embryonic development and cell proliferation. Dot plot graphic in **(a)** shows the number of E14.5 embryos from WT and DJ-1 KO mice. Dot plot graphic in **(b)** shows the brood size in *dj-1* (*djr1*.*1*/*djr1*.*2*, ΔΔ) and *glod-4* KO worms. Dot plot graphic in **(c)** shows the percentage of sperm with progressive rapid motility in WT and DJ-1 KO mice. Dot plot graphic in **(d)** shows the abortion rate (number of aborted embryos/total number of embryos) in WT and DJ-1 KO mice. Graphic in **(e)** shows the evolution of the abortion rate with age. Dot plot graphic in **(f)** shows the abortion rate for WT and DJ-1 KO mice. Dot plot graphic in **(l)** displays embryonic lethality in WT, ΔΔ and *glod-4* worms shown as the percentage of hatched eggs to the total of laid eggs. **(g)** Growth curve of WT and DJ-1 KO HeLa cells during 144 h. Graphic in **(h)** shows the effect of knocking out DJ-1 in the growth of HeLa clones using CRISP/Cas-9. Error bars correspond to S.E.M., statistical analysis was performed using a Student’s t-test **(a-d)** 1-way ANOVA with a Bonferroni post-hoc test **(f and h)**, or a 2-way ANOVA with a Bonferroni post-hoc test **(g)**. n.s. means non-significant. *,** and *** correspond to *P*>0.05, *P*>0.01 and *P*>0.001 respectively.

It has been reported that, in HeLa cells, knocking down *DJ-1* alters the rounding force exerted when the cells round-up to prepare and perform mitosis. The generation of a rounding force during mitosis is important for the transitions between different mitotic phases. This is also crucial during embryonic development ^17^. To elucidate whether knocking down *DJ-1* also lead to a defect in mitosis and, thereby, cell proliferation, we analyzed cell proliferation in WT and *DJ-1* KO HeLa cells, generated by CRISPR/Cas9. Our results show a significant delay in the growth curve of *DJ-1* KO HeLa cells when compared to WT (Figure 1g). Interestingly, this delay varied between the eight different *DJ-1* KO clones (Figure 1h), suggesting that subtle gene-environment interactions had an important effect on this phenotype.

### Glycolic acid and (partially) D-lactate increase sperm motility and rescue both the embryonic lethality and the cell proliferation phenotypes

We then tested whether treatment with GA or DL, the products of DJ-1 and GLO could rescue the phenotypes observed in *DJ-1* KO mice and ΔΔ-*djr* and *glod-4* worms. We first analyzed the effect of 10 mM GA or DL on WT and *DJ-1* KO mouse sperm motility. Adding GA, but not DL, to cauda sperm rescued the defect in Grade A rapidly progressive sperm motility in *DJ-1* KO mice (Figure 2a) and had a positive effect on overall sperm motility (increase in the percentage of spermatozoa with Grade A+B motility) also on WT mice (Figure 2b). The positive general effect of GA on mouse sperm motility was also shown for bull and human sperm using 30 and 60 mM GA (Figure 2c), demonstrating that the effect of GA on sperm Grade A+B motility is conserved between species. Remarkably, this effect was maintained over time for at least 90 min in all species (Figures 2d-h).

**Figure 2.**
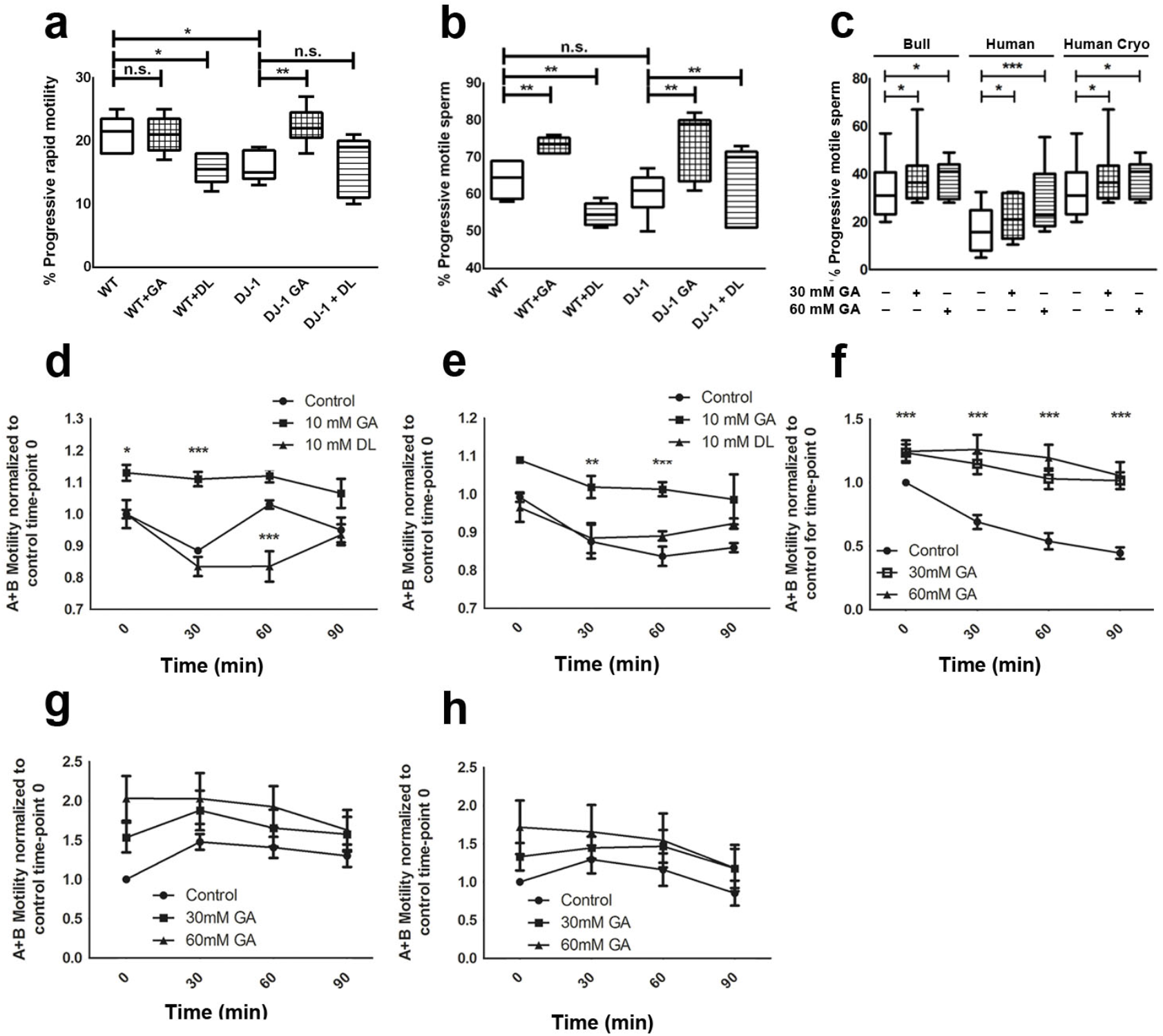
GA but not DL rescues DJ-1 dependent sperm motility deficits and has an overall enhancing effect on sperm motility. Box-and-Whisker plot graphics in **i** and **j** show the percentage of total motile **(a)** and progressive motile **(b)** sperm from WT and DJ-1 KO mice and the effect of GA and DL treatments on sperm motility. Box-and-Whisker plot graphic in **(a)** shows the percentage of progressively motile cryopreserved bull and human (fresh and cryopreserved) sperm 30 min after incubation in vehicle, 30 and 60 mM of GA. Graphics in **(d** and **f)** show the midterm effect of GA and DL on sperm motility in WT and DJ-1 mice. Graphics in **(f**, **g**, and **h)** show the midterm effect of GA on the motility of defrosted bull, fresh human and defrosted human sperm respectively. 2-way ANOVA: ctrl. vs. 30 or 60 mM GA in bull sperm (p<0.001), ctrl. vs. 60 mM GA in human sperm (p<0.05), ctrl. vs. 30 mM GA (n.s.), ctrl. vs. 30 (n.s.) or 60 mM GA in cryopreserved human (p<0.05).. Error bars correspond to S.E.M., statistical analysis was performed using a 1-way ANOVA with a Bonferroni post-hoc test **(a-c)**, or a 2-way ANOVA with a Bonferroni post-hoc test **(d-h)**. n.s. means non-significant. *,** and *** correspond to *P*>0.05, *P*>0.01 and *P*>0.001 respectively.

The effect of GA and DL on embryonic development was tested on ΔΔ-*djr* and *glod-4* worms and their effect on cell proliferation in HeLa cells. Both GA and DL treatments rescued the embryonic lethality phenotype in worms (Figure 3a) whereas only the addition of GA, but not DL, rescued the cell proliferation phenotype of HeLa cells (Figure 3b). Altogether, these results show that loss of *DJ-1* affects progressive, sperm motility, embryonic development, and cell proliferation, and that these cellular defects can be rescued by the addition of GA and, in some phenotypes, DL.

**Figure 3.**
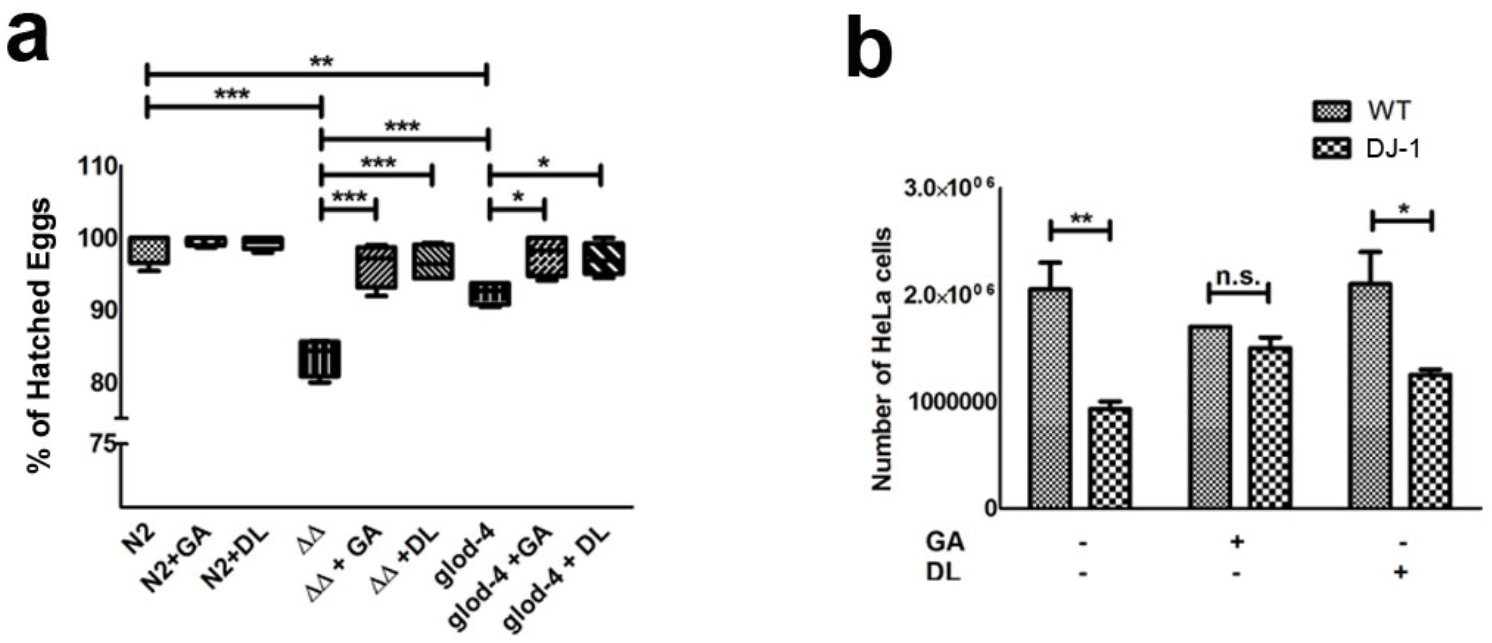
GA and DL rescue DJ-1 and glod-4 dependent embryonic lethality in C. elegans and GA but not DL rescues DJ-1 dependent cell proliferation defects in HeLa cells. Box-and-Whisker plot graphic in **(a)** display embryonic lethality in WT, ΔΔ and *glod-4* worms shown as the percentage of hatched eggs to the total of laid eggs and the effect of GA and DL treatment on embryonic lethality. **(b)** Effect of DJ-1 loss of function and the effect of GA and DL treatment on HeLa growth after 144 h. Error bars correspond to S.E.M., statistical analysis was performed using a 1-way ANOVA with a Bonferroni post-hoc test **(a)** or a Student’s t-test **(b)**.

### Glycolic acid and D-lactate decrease intracellular calcium and glycolic acid, but not D-lactate, increases mitochondrial energy production

Based on our previous results showing that GA and DL supported mitochondrial membrane potential, we first analyzed whether the effect on sperm motility was mediated by an increase in mitochondrial activity. The mid piece of the sperm flagellum contains mitochondria, which together with glycolysis has a role in energy production ^18^ ensuring reproductive success. Sperm progressive motility has been correlated with mitochondrial membrane potential (ψ_m_) and consequently with mitochondrial functionality: an increase in ψ_m_ is directly proportional to an increase in progressive motility and oxygen consumption in human sperm^19^. In clinical studies, mitochondrial function is generally monitored using cationic fluorescent probes such as MitoTracker or JC-1^20^. Using JC-1, Paoli et al. were able to demonstrate that both human sperm motility and viability are associated with ψ_m_^21^. The authors found that the higher the fluorescence signal, the better the membrane integrity of the mitochondria and a higher motility. Therefore, we stained human spermatozoa with JC-1 to analyze the effect of GA on the ψ_m_. The percentage of sperm with active mitochondria between GA-treated and control samples did not differ at the beginning. If something, treatment with 60 mM GA caused a very small but significant decrease in the percentage of bull and human sperm with higher fluorescence values (Figure 4a-c). Thus, an improvement of the mitochondrial membrane integrity does not seem to underlie the increase in progressive motility after GA treatment. We therefore looked for other factors that could improve sperm motility.

**Figure 4.**
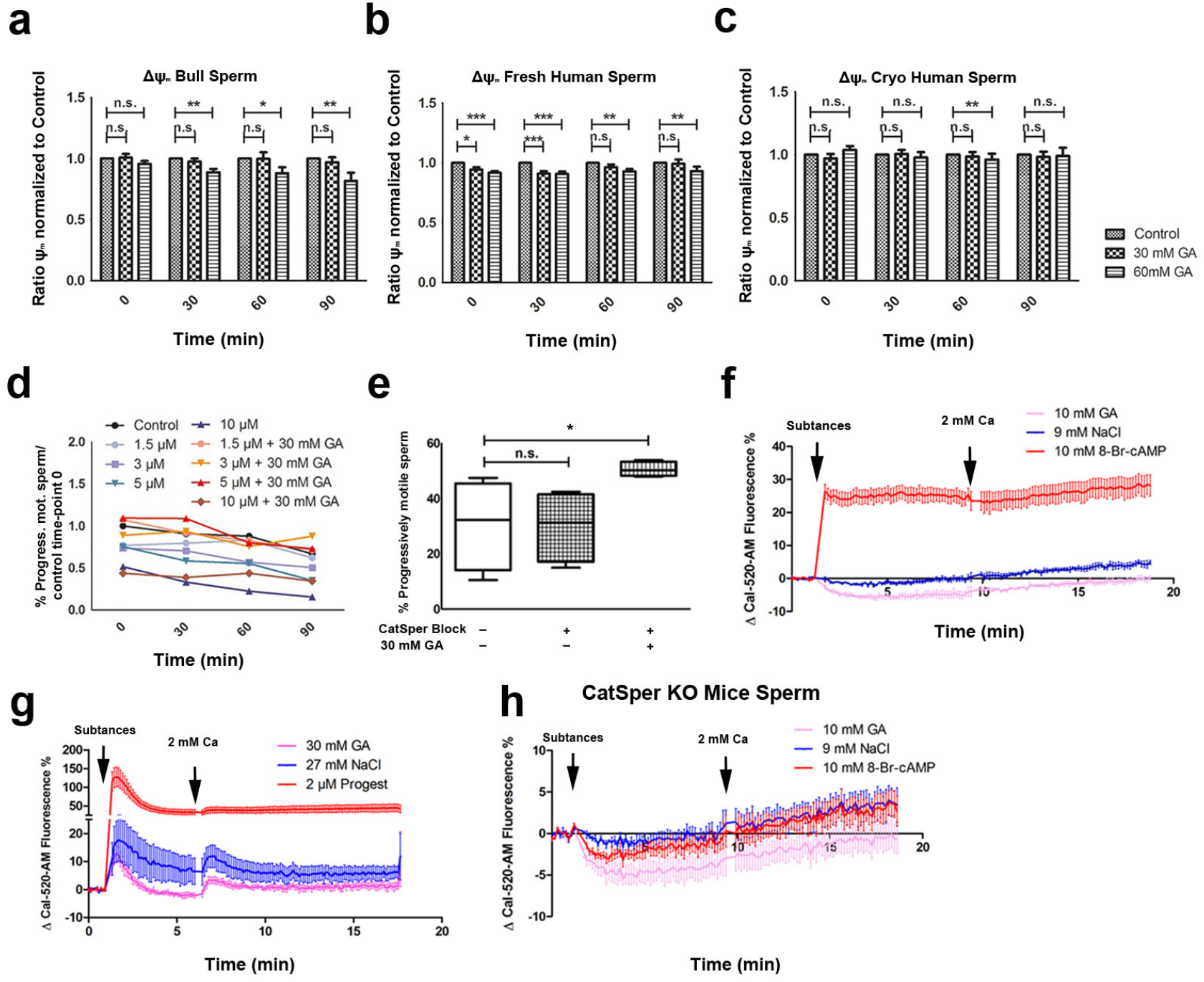
GA slightly decreases mitochondrial membrane potential and calcium content in sperm cells but rescues the effect on motility of NNC55-0396 a CatSper inhibitor. Bar graphs in **(a, b** and **c)** show the effect of GA treatment on ψ_m_ on bull, fresh human and frozen human sperm respectively, measured as the percentage of sperm with JC-1 olig./JC-1 mon. values above a certain threshold (set as a gate in the FACS analysis). Line graphic in **(d)** shows the effect of blocking CatSper with 1.5, 3, 5, or 10 µM NNC55-0396 and the co-treatment with GA on human progressive sperm motility. Statistical analysis was performed using 2-way ANOVA: Ctrl. vs. 3µM (p<0.05), ctrl. vs. 5 µM (p<0.05), ctrl. vs. 3µM + GA (n.s.), 3 µM vs. 3 µM GA (p<0.05), ctrl. vs. 5 µM+GA (n.s.), 5 µM vs. 5 µM+GA (p<0.05), ctrl. vs. 10 µM (p<0.05), 10 µM vs. 10 µM +GA (n.s.). Box-and-Whisker plot graphic in **(e)** shows the effect of NNC55-0396 on bull sperm progressive motility and the enhancing effect of GA. Graphics in **(f)** to **(h)** show variations in the Cal-520-AM fluorescence signal upon treatment with 30mM GA, 27mM NaCl and 2µM progesterone in the absence of Ca^2+^ and after the addition of Ca^2+^ in human sperm **(g)** and upon treatment with 10mM GA, 9mM NaCl and 10mM 8-Br-cAMP in the absence of Ca^2+^ and after the addition of 2mM Ca^2+^ in sperm coming from WT **(f)** and CatSper KO **(h)** mice. Error bars correspond to S.E.M.. Statistical analysis was performed using a Student’s t-test **(a-c)**, 1-way ANOVA with a post-hoc Bonferroni’s multiple comparison test **(e)** or a 2-way ANOVA **(d**,**f-h)** were used. n.s. means non-significant. *,** and *** correspond to *P*>0.05, *P*>0.01 and *P*>0.001 respectively.

Sperm motility also relies on a tight regulation of intracellular Ca^2+^ levels ^22^. The principal Ca^2+^ channel in mammalian sperm is the CatSper channel, specially in humans. Human CatSper is activated upon depolarization of the membrane potential and alkalization of the intracellular pH, but also by extracellular ligands, acting as a polymodal, chemosensory Ca^2+^ channel that plays a vital role in the regulation of sperm hyperactivation ^23-25^. In the absence of CatSper, sperm do not hyperactivate and cannot fertilize the oocyte, resulting in male infertility^25^. In addition, it has been suggested that the mobilization of intracellular Ca^2+^ stores through inositol 3-phosphate receptors (InsP3Rs) ^26,27^ and Ryanodine receptors (especially RyR 1 and 2) ^28^ located at the acrosome and the sperm neck, respectively, also control sperm function. To test whether the effect of GA on sperm motility is dependent on Ca^2+^ influx through CatSper, we first used a pharmacological approach. Human and bull sperm were treated with NNC55-0396, which inhibits CatSper channel function already at low µM concentrations ^24,29^. In human sperm, treatment with 1.5, 3, 5 or 10 µM NNC55-0396 reduced sperm motility and co-treatment with GA rescued this effect in all cases except for the 10 µM concentration (Figure 4d). In contrast and surprisingly, NNC55-0396 had no negative effect on bull sperm motility and GA increased sperm motility in its presence (Figure 4e). The positive effect of GA on human sperm motility in the presence of the blocker suggests that the effect of GA was independent of CatSper. We then tested the effect of GA on intracellular calcium. Sperm cells were loaded with Cal520-AM and the changes in fluorescence were recorded using a plate reader before and after the addition of GA. Addition of GA to mouse sperm did not evoke an increase in [Ca^2+^]_i_ as observed in WT mice exposed to 8-Br-cAMP, which directly activates CatSper (Figure 4f). If something, GA induced a non-specific but significant reduction in [Ca^2+^]_i_ when compared to control sperm. This same effect could also be observed in the absence of CatSper in sperm from *CatSper* KO mice (Figure 4h). Again suggesting that this effect was independent of CatSper. Similar to mouse sperm, in human sperm, addition of GA evoked a non-specific but significant reduction in [Ca^2+^]_i_ (Figure 4g). Thus, whereas it is unclear how the small but significant reduction in [Ca^2+^]_i_ would lead to an increase in sperm motility (discussed in detail below), the increase in sperm motility caused by GA in human and mouse sperm cannot be explained by an increase in [Ca^2+^]_i_.

The next logical step would have been to measure energy production in sperm cells. However, this is a complicated assay and it is unclear how informative the results from such an assay would have been. This is so because, even if GA increases energy production in sperm, it also increases sperm motility, which is associated with a higher energy consumption. Therefore, we tested the effect of GA and DL on energy production in HeLa cells. Our previous results had shown that *DJ-1* loss of function altered mitochondrial membrane potential (increased in worms and decreased in HeLa cells) and GA and DL were able to rescue this phenotype. However, as in the case of sperm cells, these results gave no real information as to whether these substances have a positive effect on energy production. In HeLa cells NAD(P)H levels can be measured using NAD(P)H auto fluorescence^16^ (Figure 5a). As HeLa cells do not move, one would not expect any alterations in energy consumption in these cells by adding GA or DL as in the case of spermatozoa. Our results show that in these cells, GA but not DL, increased NAD(P)H levels (see Figure 5b).

**Figure 5.**
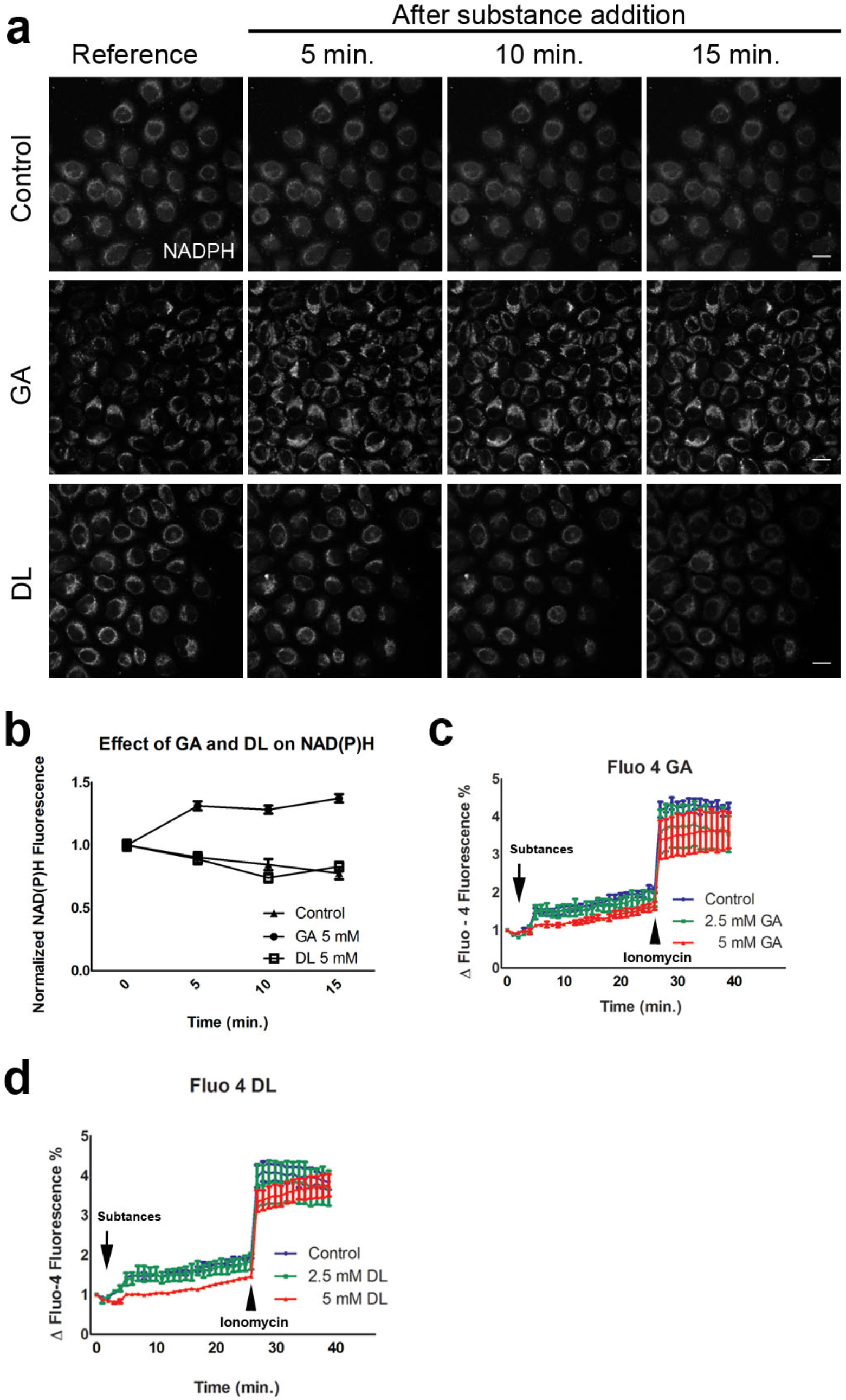
GA but not DL increases NAD(P)H fluorescence in HeLa cells and both substances decrease the concentration of intracellular calcium. Sequential microscope photos in **(a)** show the effect of GA and DL on NAD(P)H autofluorescence. Scale bar: 20 µM. As shown in graphic in **(b)**, GA but not DL treatment increased NAD(P)H fluorescence when compared to control. 2-way ANOVA: Ctrl. vs. GA (p<0.001), ctrl. vs. DL (n.s.). Error bars correspond to S.E.M. Statistical analysis was performed using a 2-way ANOVA **(b)**, a 1-way ANOVA with **(c)** or without **(d)** post-hoc Dunnett’s test. n.s. means non-significant. *,** and *** correspond to *P*>0.05, *P*>0.01 and *P*>0.001 respectively.

Finally, based on the results obtained on [Ca^2+^]_i_ in sperm, we tested the effect of GA and DL on the [Ca^2+^]_i_ in HeLa cells. We charged HeLa cells with the Ca^2+^ fluorescent marker Fluo-4 AM and analyzed the changes in [Ca^2+^]_i_ caused by the addition of GA, DL or vehicle. In these cells, the addition of 5 mM (or higher) but not 2,5 mM of GA or DL resulted in a significant decrease of [Ca^2+^]_i_ when compared to the osmotic control (NaCl) (Figure 5c and d).

## Discussion

Alterations in sperm motility and/or embryonic development lead to a reduction in fertility. In this study we show that knocking down DJ-1 reduced the litter size in mice and knocking down *djr-1*.*1/djr-1*.*2* or *glod-4* reduced the brood size in worms. In mice, the litter size reduction was associated with a reduction in sperm motility and surprisingly to an age-dependent increase in the abortion rate, which had not been previously described. These findings were reproduced in *C. elegans*. In this model organism, the brood size reduction was also associated to an increase in embryonic lethality.

Abortion and embryonic lethality are associated to alterations in embryonic development. During embryonic development, cells constantly divide and migrate. The migration process is associated with different kinds of forces exerted on the cells. This forces also affect embryonic cell division^30^. A recent article showed that DJ-1 is involved in mitotic rounding and the absence of DJ-1 leads to a reduction in the force exerted during mitotic rounding in the initial phases of mitosis^17^. To divide, most animal cells drastically change shape and round up against extracellular confinement. This process is even more important during embryonic development, as embryonic cells need to be able to counteract external migrating forces to divide properly. Therefore, we analysed whether knocking down *DJ-1* in HeLa cells would lead to alterations in cell proliferation. Indeed, our results showed that knocking-down *DJ-1* in HeLa cells leads to a reduction in cell proliferation.

In general and under controlled environmental conditions, the mutation penetrance of knocking down *DJ-1* in mice and HeLa cells or its orthologs or *glod-4* in worms suggest that gene-enviroment interactions play an important role in the development of these phenotypes. Knocking-down *DJ-1* in mice or in HeLa cells induced an overall mild negative effect (15-20% difference to wild-type for most of the values measured) and, in the case of abortion, appeared only in older animals. In HeLa cells, the overall effect on cell proliferation was mild even with some clones showing no clear phenotype. *djr-1*.*1/djr-1*.*2* and *glod-4* KO worms only exhibited a mild impairment in embryonic development. In worms the effect of these mutations on sperm motility remains to be further analyzed.

These results correlate well with previous published studies that analysed the phenotype of *DJ-1* KO mice showing no neurodegeneration or any other clear phenotype apart from a mild alteration in the mitochondrial function and an increase in glutathione peroxidase activity^31^. They also correlate well with our previous results showing that knocking out *djr1*.*1/djr1*.*2, glod-*4, or all three genes together in *C. elegans* did not have any obvious effect on the life cycle of the worm under normal environmental conditions, apart from increasing the mitochondrial membrane potential^15^. Interestingly, under stress conditions, the penetrance dramatically increased. The lack of these genes in worms caused: i) a decreased tolerance to desiccation and posterior rehydration with mortality rates up to 80% when compared to N2 worms and associated with the disruption of the mitochondrial network^15^ and ii) a decreased tolerance to PQ exposure, also increasing mortality rates. Thus, further underlining the importance of gene-environment interactions. These phenotypes were stronger in the triple mutant compared to *ΔΔdjr1*.*1/djr1*.*2* or *glod-4* double or single mutants, suggesting that the remaining gene/-s were able to partially compensate the loss of the other gene/-s. Desiccation/rehydration of worms can be observed when the soil dries out and loses humidity. Under these conditions, worms protected themselves by up-regulating the expression of *djr-1*.*1, djr1-2*, and *glod-4* (among other genes) in a process called preconditioning that takes place when the worms sense a reduction in room humidity ^32^. This up-regulation seems to be important to prepare for rehydratation, where a boost in the production of glyoxal and methylglyxoal can be expected and the up-regulation of these genes should lead to an increase in GA and DL levels. In that study, we also showed that feeding GA and DL to the mutant worms was able to rescue these phenotypes. Thus, suggesting that an increase of GA and DL levels is part of the survival strategy of *C. elegans* in dry environments and is required to overcome and protect against the stress induced by this metabolic stop/start, a process that could be compared to ischemia/reperfusion damage. A similar protective effect was observed upon exposure to mitochondrial toxins like PQ, that increased mortality rates in mutated worms compared to N2, and was rescued by the addition of GA and, to a lesser extent, DL. These results were also reproduced in murine dopaminergic neurons *in vitro*.

In contrast to *C. elegans*, no up-regulation of *DJ-1* or *GLOD-4* levels has been observed in mammals. In the literature, only a mild up-regulation of *GLOD-4* expression in a mouse model of Alzheimer’s disease has been described^33^. Despite the inability to regulate this genes, our results show that high concentrations of GA and DL still have neuroprotective properties in mammals. In a recently published study, we showed that GA protects against ischemia in several stroke models: in an oxygen glucose deprivation (OGD) model *in vitro*, in a global cerebral ischemia mouse model and in a middle cerebral artery occlusion mouse model ^34^. We have also been able to reproduce the *in vitro* results in dopaminergic neurons in a mouse model of PD generated by exposing WT and DJ-1 KO mice to PQ. In this ongoing study, we also observed that knocking down DJ-1 in mice had no effect on the concentration of GA in serum (reference to be added). Considering that GA and DL only exert their effect when using high concentrations of GA and DL (at least orders of magnitude higher than the physiological concentrations detected in the plasma of mice, i.e. >1 mM vs. <0.1 mM), these results suggest that mammals might have lost this gene regulatory pathway during evolution and that the observed rescue and protective effects of high concentration of GA and DL is a conserved evolutionary trait in higher organisms, even if they lost the ability to up regulate the expression of these genes. Supporting this hypothesis, overexpression of *DJ-1* has been described as neuroprotective in several studies using mammalian cell lines ^35,36^.

Based on these results, we investigated whether GA and DL could rescue the phenotypes observed in sperm motility, embryonic development or cell proliferation and tried to identify the possible underlying mechanisms. Overall, our results show that GA and DL worked to a different extend depending on the phenotype. Whereas treatment with GA rescued all phenotypes and increased sperm motility even in WT sperm, treatment with DL only had a positive effect on rescuing embryonic development in worms. Our results also suggest that GA exerts its effect through i) a decrease in [Ca^2+^]_i_ and ii) an increase in NAD(P)H production, most probably mitochondrial. This does not seem to be the case for DL, that had a similar effect on [Ca^2+^]_i_ but showed no effect on NAD(P)H production, suggesting that this unique feature of GA might underlie the differences in their effect on these phenotypes. The reason for this difference needs to be further investigated. Interestingly, many neurological diseases are associated with an impaired mitochondrial function and an increased intracellular calcium, thought to play an important role in the pathophysiological process. Therefore, the effects of GA and DL on intracellular calcium and/or mitochondrial function could also explain their neuroprotective properties.

The results also suggest that these two different mechanisms do not need to take place at the same time to elicit a positive effect. It seems that this would depend on the cell type and whether only one or both mechanisms are needed to exert a beneficial effect in that specific cell. For example, in the case of sperm the possible role of reducing [Ca^2+^]_i_ to enhance sperm motility is unclear. Reducing the concentration of Ca^2+^ in i-TALP buffer did not increase sperm motility (data not shown). Moreover, calcium influx through CatSper channels and the resulting increase in intracellular Ca^2+^ are essential for sperm motility ^24,37^. Accordingly, blocking CatSper channels with NNC55-0396 reduced human sperm motility. GA was able to counteract the inhibitory effect of NNC55-0396 on sperm motility up to a concentrations of 5 µM. Thus suggesting that the effect of GA is independent of the regulation of [Ca^2+^]_i._ This was also the case in bovine sperm, even if, surprisingly, NNC55-0396 did not show any effect on bovine sperm motility. A literature search did not find any studies investigating the effect of NNC55-0396 on bovine sperm motility, so it could be that NNC55-0396 does not affect bovine sperm motility. Another argument in this direction is the lack of other sources of Ca^2+^ in sperm cells. Mammalian sperm cells only contain a reminiscent of the ER and the presence of other Ca^2+^ channels (e.g. ER-related), apart from CatSper channels, is still a matter of debate ^26,27,37-39^. Therefore, it seems more plausible that the positive effect of GA on sperm motility is due to an increase in energy production. In this regard, JC-1 staining did not have any clear effect on mitochondrial membrane potential. The next logical step would have been to compare ATP or NAD(P)H production between control and GA treated sperm. These experiments are complicated to perform and we believe that, in the specific case of GA, that also enhances sperm motility, it would have been difficult to extract a conclusion from such measurements. This is so because GA also increases sperm motility, which would also increase energy consumption. This has been shown in capacitated sperm. Capacitation dramatically increases sperm motility and energy consumption which is then partially compensated by a boost of energy production from glycolysis and mitochondrial oxidative phosphorylation (OXPHOS), the two major metabolic pathways producing ATP in sperm (reviewed in ^40^). The fact that only GA but not DL can increase sperm motility also suggests that increases in mitochondrial energy production would be more important than Ca^2+^ regulation in exerting this effect. This assumption is based on the data obtained with HeLa cells. As mentioned above, our results show that GA reduces [Ca^2+^]_i_ and increases NAD(P)H production in this cell type. The distribution pattern of the NAD(P)H signal obtained with the confocal microscope, similar to the pattern of the mitochondrial network, suggests that NAD(P)H had a mitochondrial origin. On the other hand, DL only decreased intracellular Ca^2+^ having no effect on energy production.

A review of the literature shows how [Ca^2+^]_i_ and energy availability also play an important role in embryonic development, mitosis and how alterations in these processes can have severe consequences and result in several diseases ^41-43^. It is therefore not surprising that, by modulating [Ca^2+^]_i_ and enhancing energy production, GA and DL have such an effect not only in these cellular functions but also rescuing pathological conditions such as neurological diseases. Further studies are necessary to understand how reducing [Ca^2+^]_i_ and increasing energy production rescue the effect of knocking down *DJ-1* on mitosis and embryonic development and whether this would also be beneficial in other diseases or in aging.

Overall the results of this study could have far-reaching implications, not only in the field of neurology, reproduction and embryonic development, but also in other diseases or in aging, where alterations in calcium homeostasis and mitochondrial dysfunction or insufficient energy production are common underlying pathophysiological features.

## Author’s contribution list

FP-M conceived and designed the study. FP-M planned and perform the experiments in mice. SB, YD, RN, IA, MT and FP-M designed and performed the sperm motility experiments. MB designed and performed the calcium measurements in sperm. IP, IR-A, YD, AAH and FP-M planned and performed the HeLa experiments. FP-M planned and performed C. elegans embryonic lethality experiments. IR-A, YD, SB, MT, IP, HvH, CK, RB, RO, HP-S, MD, CGS, MB, AAH and FP-M analyzed the data. MB, IP, SB, IR-A, MT, HP-S and FP-M wrote the manuscript and all other authors critically corrected and reviewed the manuscript.

## Acknowledgements

This work was funded by the Deutsche Forschungsgemeinschaft (DFG, German Research Foundation) under Germany’s Excellence Strategy within the framework of the Munich Cluster for Systems Neurology (EXC 2145 SyNergy – ID 390857198). We would like to thank Dr. Viktoria Ruf for her help with the fluorescence plate reader. *C. elegans* mutant lines were generated and kindly provided by Prof. Teymuras Kurzchalia and Dr. Cihan Erkut. Prof. Teymuras Kurzchalia was also of great help critically revising the manuscript. CRISP/Cas-9 modified HeLa Park-7^-/-^ clones were kindly provided by Dr. Martin Steward from the MIT. Dr. Ünal Coskun was of great help critically reviewing the manuscript and proposing additional critical experiments. Thanks to Wolfgang John and Jussi Helppi from the MPI Animal Facility for their help with PARK-7 mice. We would also like to thank the animal caretakers at the Zentrum für Neuropathologie und Prionforschung, especially Dr. G. Mitteregger, Fang Zhang, Heike Jäckle and Tanja Simon for their help with the animals.

